# Monocyte to macrophage differentiation and changes in cellular redox homeostasis promote cell type-specific HIV latency reactivation

**DOI:** 10.1101/2024.02.12.579955

**Authors:** Alexandra Blanco, Robert A. Coronado, Neha Arun, Kelly Ma, Roy D. Dar, Collin Kieffer

## Abstract

Human Immunodeficiency Virus (HIV) latency regulation in monocytes and macrophages can vary according to signals directing differentiation, polarization, and function. To investigate these processes, we generated an HIV latency model in THP-1 monocytes and showed differential levels of HIV reactivation among clonal populations. Monocyte-to-macrophage differentiation of HIV-infected primary human CD14+ and THP-1 cells induced HIV reactivation and showed that virus production increased concomitant with macrophage differentiation. We applied the HIV-infected THP-1 monocyte-to-macrophage (MLat) model to assess the biological mechanisms regulating HIV latency dynamics during monocyte-to-macrophage differentiation. We pinpointed PKC signaling pathway activation and Cyclin T1 upregulation as inherent differentiation mechanisms that regulate HIV latency reactivation. Macrophage polarization regulated latency, revealing pro-inflammatory M1 macrophages suppressed HIV reactivation while M2 macrophages promoted HIV reactivation. Because macrophages rely on reactive-oxygen species (ROS) to exert numerous cellular functions, we disrupted redox pathways and discovered that inhibitors of the thioredoxin (Trx) system acted as latency promoting agents (LPAs) in T-cells and monocytes, but opposingly acted as latency reversing agents (LRAs) in macrophages. We explored this mechanism with Auranofin, a clinical candidate for reducing HIV reservoirs, and demonstrated Trx reductase (TrxR) inhibition led to ROS induced NF-κB activity, which promoted HIV reactivation in macrophages, but not in T-cells and monocytes. Collectively, cell type-specific differences in HIV latency regulation could pose a barrier to HIV eradication strategies.

## Introduction

Despite the success of antiretroviral therapy (ART) at suppressing viral replication and improving the quality of life of people living with Human Immunodeficiency Virus (PLWH; HIV), HIV remains a major public health challenge. Upon cessation of ART, a pool of cells harboring non-productive or latent infections can reactivate to produce virus and lead to rebound in the levels of viremia (*1*). These cells form stable and long-lived latent reservoirs throughout the body and are a major hurdle to eradicating HIV (*2*). Current cure-based efforts are largely focused on targeting the latent reservoir of CD4^+^ T-cells, as these cells are the primary target of HIV infection and form the largest reservoir in the body (*3*, *4*). However, the latent HIV reservoir is composed of multiple cell types that can exhibit unique regulation of HIV gene expression (*5*, *6*).

Myeloid cells, such as monocytes and macrophages, constitute an important viral reservoir that persists in PLWH (*7*). Monocytes circulate in the bloodstream, can migrate into tissues, and differentiate into monocyte-derived-macrophages (MDMs) in response to biological cues. This property makes monocytes attractive candidates for seeding tissue reservoirs and trafficking virus between them. It is currently unknown how the transition of latently infected monocytes into MDMs can impact the stability of latent HIV. For example, it is possible that the differentiation process triggers the expression of host factors required for transcription of full-length HIV, such as the positive transcription elongation factor b (P-TEFb) or Cyclin T1 (CycT1) (*8*, *9*). Monocytes express low levels of CycT1, but during monocyte-to-macrophage differentiation, CycT1 is rapidly induced to high levels (*10*, *11*). This mechanism inherent to differentiation may trigger HIV reactivation in an MDM. In tissues, MDMs are highly plastic cells that respond to changes in their local cytokine environment and differentiate into specialized phenotypes with defined biological functions, such as M1 pro-inflammatory and M2 anti-inflammatory macrophages. Many of the cues that direct biological function and MDM differentiation, such as highly pro-inflammatory microenvironments, can trigger HIV reactivation from latency in other cell types to seed new reservoirs (*12*). Additionally, many of these anatomical reservoirs can be resistant to ART and constitute sanctuaries where target cells remain unprotected from infection (*13*, *14*). These properties, along with recent evidence indicating MDMs contribute to persistent latent reservoirs in PLWH (*15*), highlight the importance of understanding the regulation of HIV latency within monocytes and MDMs, and characterizing their contributions to latent reservoirs.

Two primary strategies are under development to combat the latent HIV reservoir. “Shock and Kill”, uses latency reversal agents (LRAs) in combination with ART to drive latently-infected cells into a productively replicating state, where they will die from viral cytopathic effects or go through immune clearance (*16*). Alternatively, the “Block and Lock” strategy uses latency promoting agents (LPAs) to force integrated HIV provirus into a deep latent state where it can no longer reactivate spontaneously (*17*). Successful application of these strategies would remove the need for lifelong ART in PLWH. However, as additional cellular reservoirs are considered, eradication strategies become more complicated. For example, HIV reactivation or suppression agents may behave in a cell-type specific manner, challenging any clinical feasibility.

A recent study discovered that certain inhibitors of the thioredoxin system behave as LPAs in T-cells (*18*). The thioredoxin system is a major component of the mammalian antioxidant defense, responsible for maintaining redox balance and scavenging reactive oxygen species (ROS), and is composed of thioredoxin (Trx), thioredoxin reductase (TrxR), and nicotinamide adenine dinucleotide phosphate (NADPH) (*19*). TrxR is a NADPH-dependent enzyme that catalyzes the reduction of oxidized Trx by electron transfer (*20*). Upon reduction by TrxR, Trx regulates the oxidation/reduction status of many proteins by disulfide bond reduction, thereby affecting protein function (*21*). The thioredoxin system is an attractive therapeutic target given its impact on HIV gene expression and latency regulation (*22–24*). Auranofin, an inhibitor of TrxR, is a promising clinical candidate that was shown to reduce total viral DNA in blood samples from humans and macaques in combination with ART (*25*, *26*). Given these promising results, it is important to evaluate the effects of modulating the thioredoxin system in multiple cell types forming the latent reservoir.

Here we developed a monocyte-to-macrophage model of HIV latency which revealed differential latency regulation in monocytes and MDMs. We showed that upon MDM differentiation, latent provirus switched to a persistent, actively replicating state, representing a risk to viral dissemination in tissues. We evaluated latency regulation dynamics in MDMs across different polarization phenotypes and highlighted multiple mechanisms contributing to latency reversal during differentiation. Disruption of redox homeostasis revealed differential cellular responses promoting latency in T-cells and monocytes and reversing latency in MDMs. Cumulatively, these studies highlight the importance of evaluating the myeloid latent reservoir at the single cell level and the potential for HIV treatment strategies to exhibit cell-type specific differences.

## Results

### Clonal populations of latently infected monocytes differentially regulate HIV latency

We generated a monocyte-to-macrophage model of HIV latency based on the well-established Jurkat T-cell latency model “JLat” and our previously-reported monocyte latency model in THP-1 cells “TLat” (*27*, *28*) (Fig. 1A). THP-1 monocytes were infected with the HIV NL4-3 ΔEnv EGFP replication incompetent vector at a MOI<1 (Fig. S1A). One week later, the resulting cell population was sorted using fluorescence-activated cell sorting (FACS) to separate GFP+ cells actively expressing HIV genes from GFP^-^ cells containing a mixed population of uninfected and latently infected cells. After one week, the GFP^-^ population was treated with tumor necrosis factor alpha (TNF-*α*), a LRA and activator of the nuclear factor kappa B (NF-κB) pathway, for 24h to activate HIV transcription. Single cells expressing GFP were sorted into individual wells of 96-well plates and expanded for 3-4 weeks to generate an isoclonal TLat library of latently infected monocytes. To assess the capacity of monocytes to reactivate from latency, a total of 67 individual clonal populations were treated with TNF-*α* for 24h, and GFP expression was quantified via flow cytometry (Fig. 1B). We divided the TLat library into three groups based on distinct reactivation profiles: low reactivation, intermediate reactivation, and high reactivation (Fig. S1B). Clones with low reactivation profiles (34 clones or ∼55% of the library) had a net change in reactivation (%GFP^+^ TNF-*α* stimulated – %GFP^+^ unstimulated) less than 1%. 25 clones (∼37% of the library) showed intermediate reactivation profiles, ranging from 1.06% to 6.98% (Fig. 1C). Clones with high reactivation profiles (5 clones or ∼7% of the library) had reactivation percentages that ranged from 41.2% to 71.4%. These results indicate that monocytes retain differences in their ability to respond to LRAs and activate transcription of latent HIV, likely influenced by integration site differences and local chromatin state (*29*). We selected four clones with intermediate reactivation profile (1C2, 2C4, 3B9, and 2C8) and five clones with high reactivation profile (2E4, 3D3, 1D5, 3B2, 4G10-1) for further characterization.

**Figure 1.**
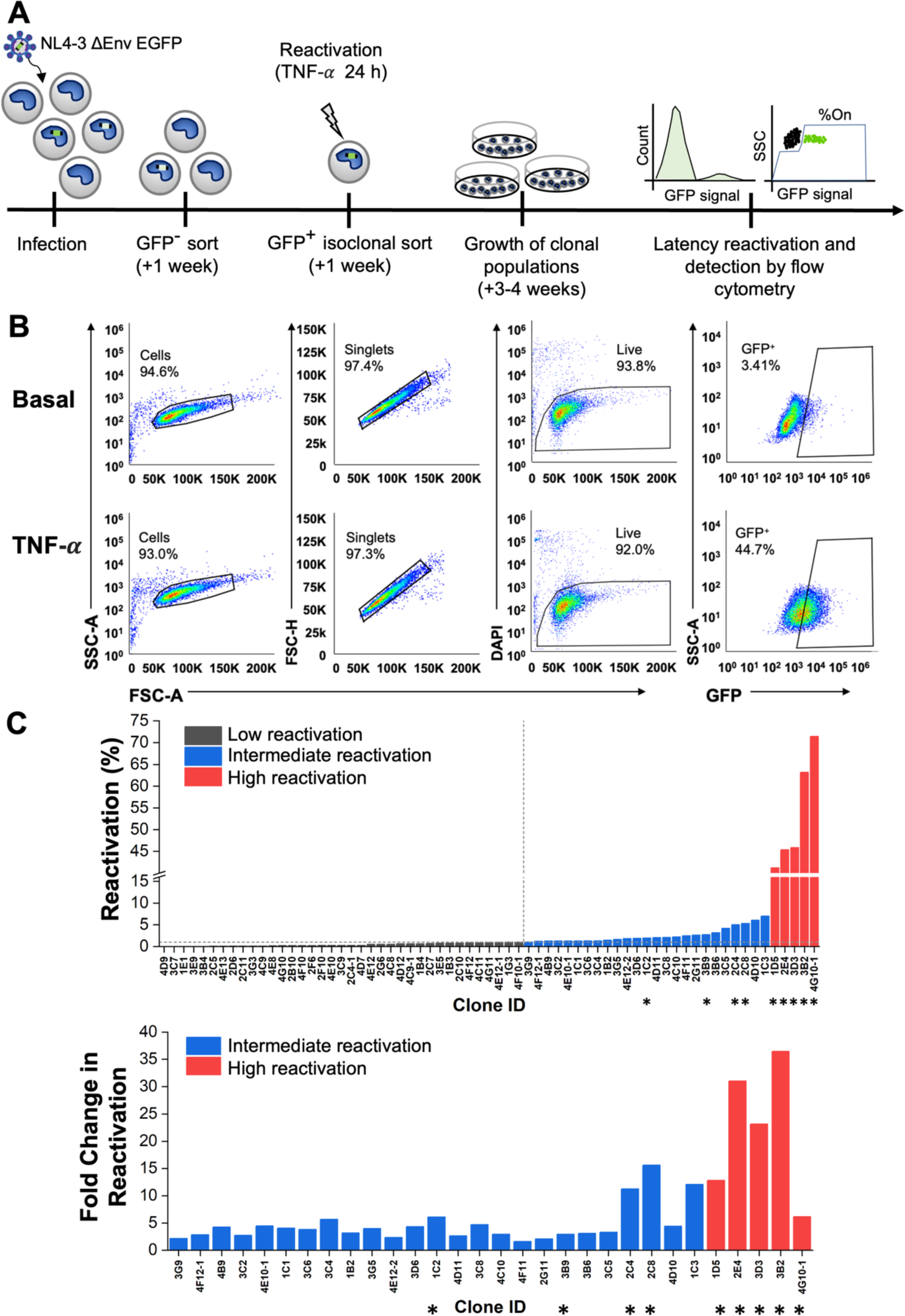
Generation of a monocyte model of HIV-1 latency. (**A**) THP-1 monocytes were infected with HIV-1 NL4-3 ΔEnv EGFP. After one week, GFP- cells were sorted by FACS and stimulated with TNF-*α* for 24 h the following week. GFP+ cells were individually sorted and allowed to relax back into a latent state to generate clonal TLat cell lines. Clonal cell lines were analyzed for GFP expression after TNF-*α* stimulation. (**B**) Representative flow cytometry gating and analysis for basal (top), and TNF-*α* (bottom) stimulated conditions. Gating was set based on the uninfected THP-1 population (Fig. S1A). (**C**) HIV latency reactivation percentage of the TLat clonal library after TNF-*α* stimulation for 24h (top). Fold change in reactivation of the intermediate and high reactivation clones (bottom). Clones denoted with asterisks were selected for further analysis.

### Monocyte to macrophage differentiation triggers HIV reactivation from latency in MDMs

HIV-infected circulating monocytes pose a risk to viral dissemination, as they can enter tissues and differentiate into resident macrophages (*30*). We propose two different viral outcomes during monocyte-to-macrophage differentiation (Fig. 2A). One possible outcome holds that differentiation does not influence the regulation of HIV latency within the cell, giving rise to a newly differentiated macrophage that harbors a latent provirus. The other possible outcome hypothesizes that differentiation triggers a cascade of transcriptional changes that destabilize latency, resulting in HIV reactivation and active viral replication in the newly differentiated cell. To test this, we differentiated several clonal TLat monocyte populations to monocyte-derived macrophages (MDMs), which we termed “MLat” cells, using phorbol 12-myristate 13-acetate (PMA) (*31*). Treatment of HIV+ and HIV- monocytes with PMA generated MDMs with commonly reported characteristics in the literature, such as cell adherence to tissue culture plastic and limited cell proliferation (*32*). Following differentiation, cells were rested in PMA-free media and analyzed via flow cytometry for GFP expression (HIV reactivation) for 6 days post-differentiation, which marks one week after PMA addition. Clones with high reactivation profile revealed an increase in HIV reactivation over time after differentiation (Fig. 2B). This latency reactivation did not appear to plateau or decrease at any timepoint, indicating sustained HIV transcription after macrophage differentiation. However, HIV reactivation followed an opposite pattern in clones with intermediate reactivation profile, remaining consistently low over time. To validate our cell line model and further assess the impact of HIV reactivation during macrophage differentiation, peripheral blood mononuclear cells (PBMCs) were processed to isolate primary CD14+ monocytes (Fig. S2A). We used a replication incompetent HIV reporter vector, HIV_GKO_, in which GFP is under the control of the HIV-1 promoter in the 5′ LTR to track virus production over time (*29*). HIV_GKO_ infectivity was tested on multiple cell lines (Fig. S2B) prior to primary CD14+ monocytes, which were refractory to infection (Fig. S2C). However, these cells became permissive to infection as they initiated macrophage differentiation, agreeing with previous literature (*33*, *34*). Thus, we treated CD14+ monocytes with macrophage colony stimulating factor (M-CSF) and infected them during the differentiation process, which can take up to ∼10 days to complete in primary cells (*34*, *35*) (Fig. 2C). We found that all primary cell donors exhibited an increase in HIV reactivation over time, concurrent with differentiation towards the macrophage lineage (Fig. 2D and Fig. 2E). We observed varying degrees of infection in individual donors, as well as varying magnitudes of virus production over time, mirroring the heterogeneity in HIV reactivation observed with clonal populations of TLat/MLat cell lines reported here. These results indicate that macrophage differentiation favors HIV latency reactivation in primary cells and confirm the validity of our TLat/MLat model to further explore the biological mechanisms of HIV latency dynamics in monocytes and macrophages.

**Figure 2.**
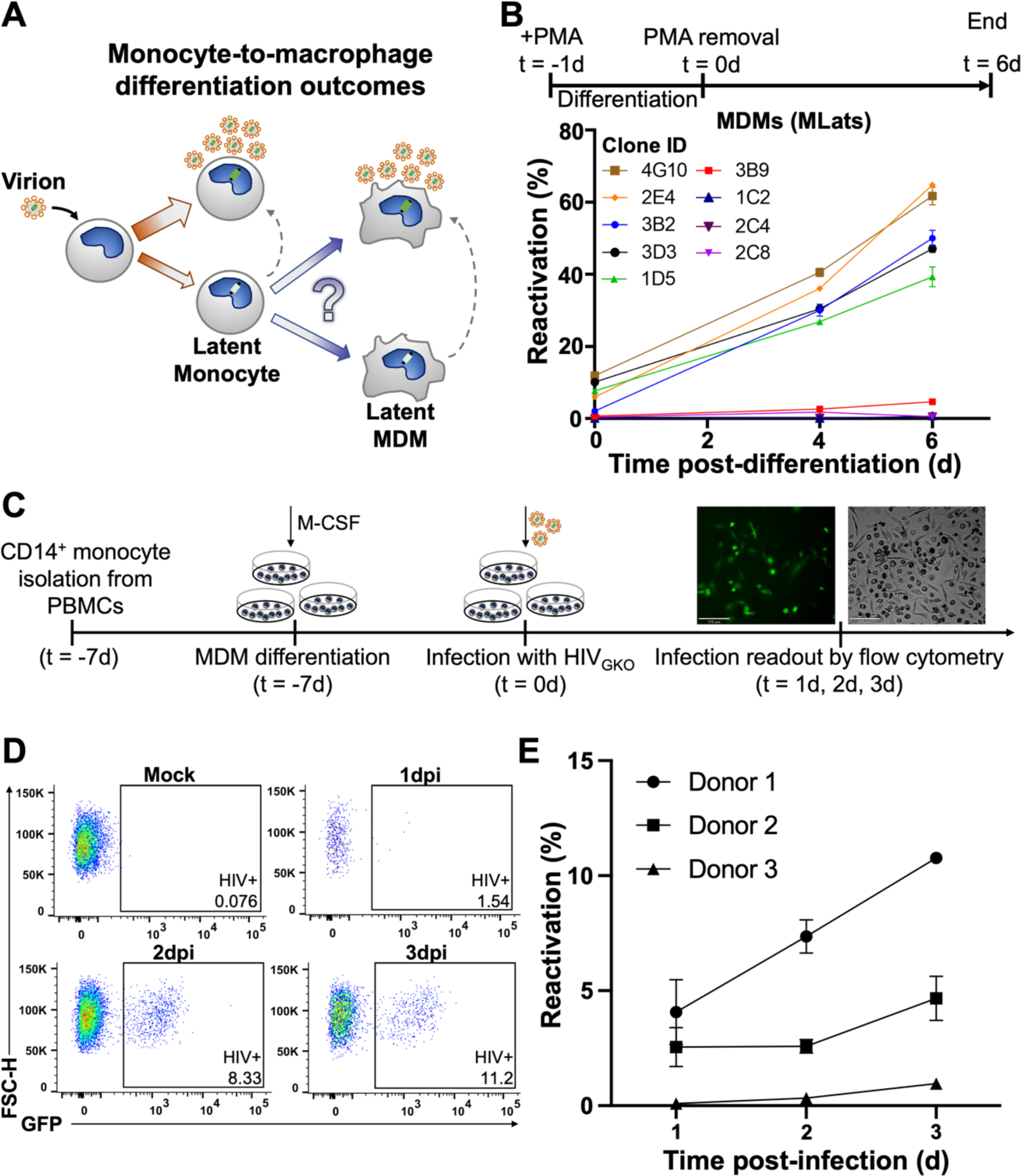
**Macrophage differentiation induces HIV latency reactivation. (A**) Schematic showing upon infection, HIV can establish a productively infected and latent state capable of spontaneous reactivation in monocytes. Differentiation of latently infected monocytes into macrophages could result in two potential viral outcomes: latency or active replication. (**B**) Latently infected monocytes (TLats) were differentiated into macrophages (MLats) with PMA for 24h. Following PMA removal, MLats were characterized at different timepoints for HIV reactivation (Top). HIV reactivation was quantified by flow cytometry based on GFP fluorescence following monocyte-to-macrophage differentiation (Bottom). Data are represented as mean of at least two independent replicates ± SEM. (**C**) CD14+ monocytes were isolated from PBMCs, macrophage differentiation was initiated with M-CSF, and cells were infected with HIV_GKO_ for 2h. HIV latency reactivation was monitored during differentiation via flow cytometry. (**D**) Representative flow cytometry gating and analysis for HIV-infected CD14+ differentiating macrophages. (**E**) HIV latency reactivation over time in CD14+ differentiating macrophages from three independent donors. Data are represented as mean of at least three independent replicates ± SEM.

### Latency reversal during monocyte-to-macrophage differentiation is attributed to differentiation mechanisms

To explore the mechanisms behind latency reversal in MDMs, we selected TLat 1D5, a representative clone with low basal expression but high fold change in reactivation after differentiation (Fig. 1C). PMA is a protein kinase C (PKC) agonist that was previously used as a latency reversal agent (LRA) *in vitro* to induce T-cell activation and the expression of latent HIV (*36*). Since PMA activates PKC and, in turn, NF-κB (*37*), it is possible that PMA stimulation could be responsible for the activation of HIV transcription during differentiation. To verify that differentiation-induced reactivation was conserved among distinct differentiation methods in our MLat model, we generated MLat cells in the presence of two alternative compounds, Vitamin D_3_ (VitD) and retinoic acid (RA), which induce macrophage differentiation, but are not PKC agonists (*38–40*). MLats were generated with VitD, and RA and VitD in combination (RA+VitD) for 72h and evaluated for HIV reactivation post-differentiation. MDMs generated by all three methods exhibited a longitudinal increase in HIV reactivation, regardless of the stimuli used (Fig. 3A). These results suggest increases in latency reactivation following monocyte-to-macrophage differentiation is independent of differentiation method, and not an exclusive result of PMA stimulation. Finally, HIV reactivation decayed after LRA removal in both T-cells (JLats) and monocytes (TLats) treated with PKC agonists (Fig. 3B & Fig. S3A), which indicates the continued increase in HIV reactivation after monocyte-to-macrophage differentiation is attributed to differentiation mechanisms.

**Figure 3.**
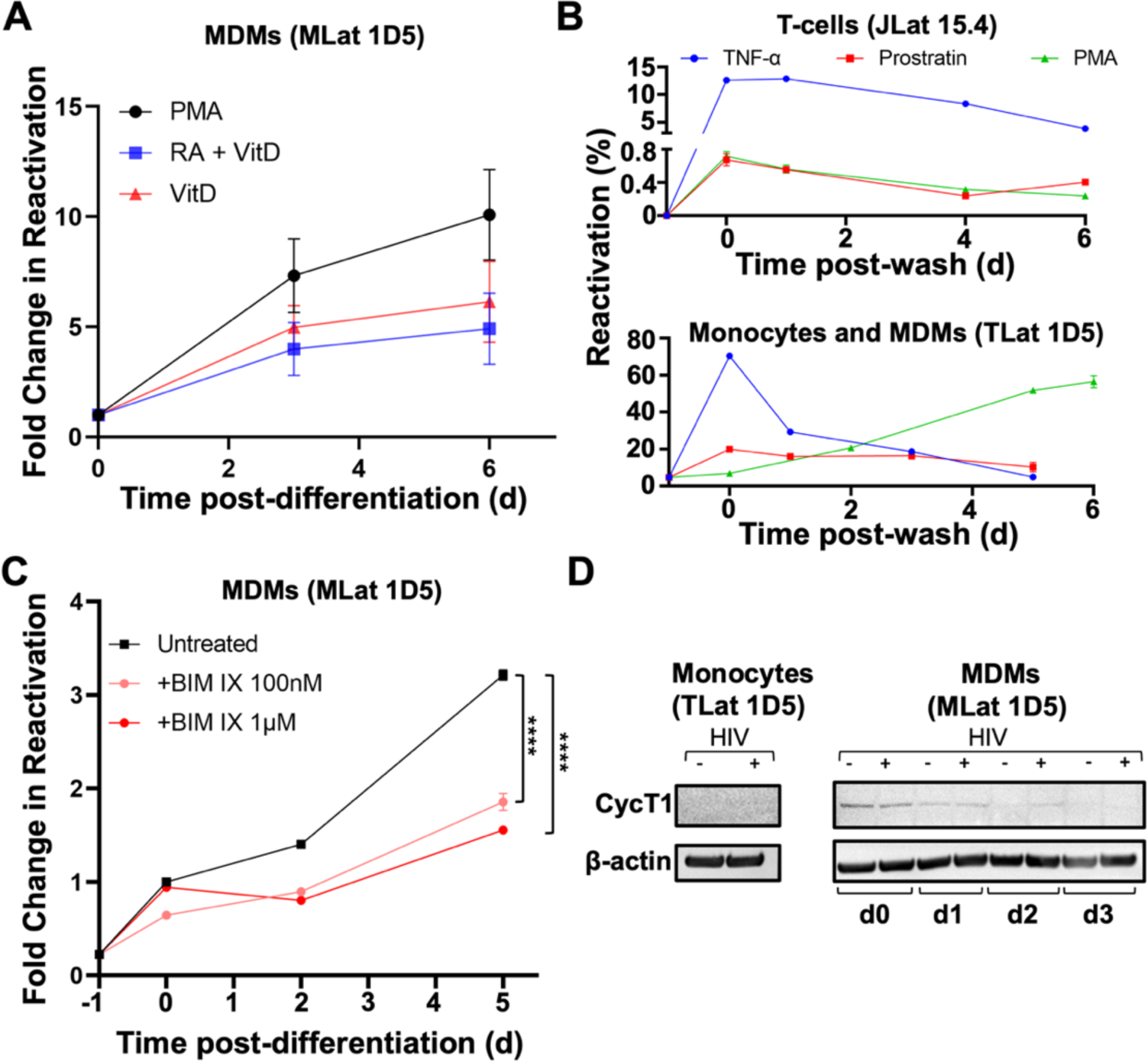
Mechanisms of HIV reactivation during macrophage differentiation. (**A**) Reactivation post-differentiation was evaluated in MLats (1D5 clone) generated by different methods (PMA, VitD, and RA+VitD). Data are represented as mean of 5 replicates from two independent experiments ± SEM. (**B**) Three LRAs (PKC agonists) TNF-***α***, Prostratin, and PMA, were added to latently infected T-cells (top) and monocytes (bottom) for 24h and removed (t=0) to evaluate HIV reactivation over time. PMA causes MDM differentiation in monocytes. Data represents the mean of at least three independent replicates ± SEM. (**C**) Latently infected monocytes were pre-treated with the non-selective PKC inhibitor BIM IX for 30min prior to differentiation into macrophages and for the duration of the experiment. HIV reactivation was quantified at different time-points following differentiation. Statistical significance was determined by performing a two-way ANOVA comparison with Dunnett correction (****: p<0.0001). Data represents the mean of three independent replicates ± SEM. (**D**) Western blot analysis of cyclin T1 (CycT1) and B-actin control in THP-1 (-HIV) and TLat 1D5 (+HIV) monocytes and MDMs.

Activation of the protein kinase C (PKC) pathway plays a role in monocyte-to-macrophage differentiation (*38*). To test the contribution of PKC activation in HIV reactivation during monocyte-to-macrophage differentiation, we pre-treated TLats with the non-selective PKC inhibitor bisindolylmaleimide IX (BIM IX or Ro 31-8220) for 30 minutes before differentiation and for the duration of PMA treatment (Fig. 3C). Importantly, BIM IX treatment did not impact macrophage morphology or alter flow cytometry forward and side scatter profiles, indicating treatment did not impact macrophage differentiation state (Fig. S3D). Following differentiation, MLats were rested in regular growth media and HIV reactivation was quantified longitudinally by flow cytometry. BIM IX treatment reduced MLat HIV reactivation by at least 1.75-fold at day 2 and 2-fold at day 5 post-differentiation, indicating that differentiation triggers the activation of PKC isoforms that are capable of stimulating HIV transcription (Fig. 3C and Fig. S3B). These results suggest that PKC is one of the molecular switches exerting a dual control on MDM differentiation and HIV reactivation.

To probe the mechanism of differentiation-induced reactivation of HIV latency in the MLat model, we evaluated CycT1 levels via western blot and found that CycT1 was absent in monocytes, but was induced after differentiation into MDMs (Fig. 3D). This induction was transient, as CycT1 levels decreased over time after differentiation. However, HIV-infected MDMs (MLats) retained higher levels of CycT1 post-differentiation compared to uninfected MDMs. HIV Gag transcript levels remained elevated post-differentiation (Fig. S3C), corresponding to sustained virus replication over time. This suggests that in MLats, elevated CycT1 levels during MDM differentiation may contribute to HIV transcription but are not required to sustain virus replication over time.

### Macrophage polarization regulates HIV reactivation capacity

Plasticity is a key characteristic of macrophages, allowing them to sense and respond to changes in their microenvironment (*30*). In response to environmental cues within tissues, macrophages acquire distinct phenotypes with defined biological functions. Toll-like receptor (TLR) ligands and type II interferon (IFN) signaling drive macrophages to undergo “classical” or M1 activation, which results in the production of pro-inflammatory cytokines, mediation of resistance to pathogens, and initiation of microbicidal mechanisms (*41*). T helper 2 (Th2) cell-associated cytokines such as interleukins 3, 4, and 10 (IL-4, IL-13, IL-10), and transforming growth factor beta (TGF-*β*) drive macrophages towards “alternate” or M2 activation, resulting in tissue remodeling, angiogenesis, mitigation of inflammation, and mediation of response to parasites (*42–44*). Macrophage polarization state was previously shown to affect susceptibility to HIV infection, replication, and latency reactivation (*45–48*). To explore whether the signaling molecules that drive polarization can affect HIV expression and latency stability, we polarized M0 THP-1 and MLat cells into M1 or M2 phenotypes (Fig. 4A). To thoroughly characterize our MLat model, we assessed cell surface marker expression (CD80, CD209, CD11b, HLA-ABC, HLA-DR, and CD4), and cytokine gene expression (IL-6, TNF-α, IL-1β, TGM2, CCL22, PPARγ) associated with each polarization phenotype, and confirmed expected macrophage polarization profiles (Fig. S4) (*49*, *50*). Following polarization, HIV latency reactivation was evaluated with flow cytometry and cell morphology changes with epifluorescence microscopy. Polarization towards an M1 phenotype decreased the amount of HIV reactivation over time, whereas polarization towards M2 resulted in the opposite effect, with M2 MLats showing increased levels of HIV reactivation (Fig. 4B).

**Figure 4.**
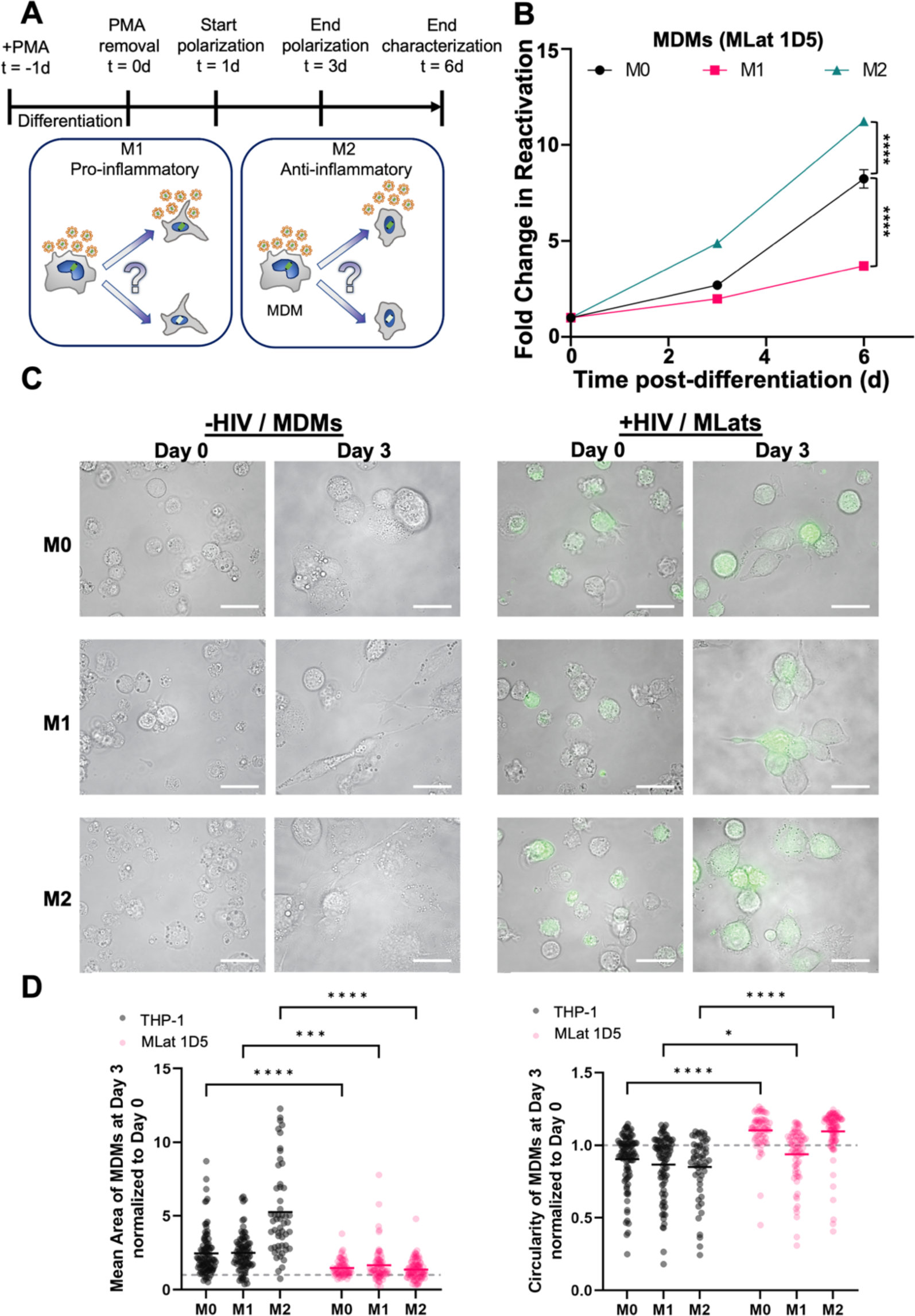
Macrophage polarization alters HIV reactivation capacity. (**A**) One day post-differentiation, MLats were polarized for 48h towards M1 or M2 phenotypes with LPS + IFN-y and IL-4, respectively (Top). Macrophage polarization towards a pro-inflammatory/M1 phenotype or an anti-inflammatory/M2 phenotype could increase or decrease viral production (Bottom). (**B**) HIV reactivation in polarized MLats was quantified by flow cytometry at different time-points post-differentiation. Data represents the mean of six replicates from two independent experiments ± SEM. Statistical significance was determined by performing a two-way ANOVA comparison with Dunnett correction (****: p<0.0001). (**C**) Fluorescence microscopy images showing levels of latency reactivation (green) and cell phenotype over time for uninfected and infected M0, M1, and M2 MDMs. All scale bars are 100 µm. (**D**) Morphological quantification of area (left) and circularity (right) of uninfected and infected M0, M1, and M2 MDMs at day 3 normalized to day 0. Circularity value ≥1 indicates a more circular shape. n ≥ 53. Statistical significance was determined by performing a two-way ANOVA comparison with Bonferroni correction (*: p<0.05, ***: p<0.001, ****: p<0.0001).

### HIV deregulates cell morphology in latently infected MDMs

Polarization of uninfected MDMs (THP-1) induced distinguishable morphological changes in M0, M1, and M2 MDMs (Fig. 4C). At day 3, uninfected M0 MDMs were more spherical, M1 MDMs acquired an elongated, spindle-like shape, and M2 MDMs exhibited a spread-out, ameboid phenotype (Fig. 4C). However, HIV-infected MDMs (MLats) showed more compact, less pronounced phenotypes for all polarization states. Quantification of cell surface area and circularity revealed that all uninfected MDMs were larger with a broader area distribution, while MLats were uniformly more circular (Fig. 4D). This was most pronounced in the M2 phenotype. Additionally, HIV-infected MDMs exhibited an overall reduction and altered distribution of cell surface marker expression when compared to uninfected MDMs for all three phenotypes studied (Fig. S4), indicating that HIV infection can deregulate the macrophage phenotype and potentially disrupt biological function. Cumulatively, these results highlight the influence that microenvironmental stimuli can have on HIV latency reactivation in macrophages and their potential contribution to viral spread.

### Inhibition of the thioredoxin system promotes latency in T-cells and monocytes but causes latency reversal in MDMs

To identify a potential biological mechanism behind the observed increase in latency reactivation following monocyte-to-macrophage differentiation, we performed targeted chemical perturbations using small molecule inhibitors of proteins that control oxidation-reduction (redox) homeostasis, which have been reported to act as latency promoting agents (LPAs) in latently infected T-cells (JLats) (*18*). Macrophages are highly sensitive to changes in cellular redox status and rely on ROS to exert their antimicrobial functions (*51*, *52*). Given the influence that redox state can have on the expression of HIV, we aimed to test whether latency regulatory dynamics are shared between different cell types and whether cellular oxidation-reduction status can influence HIV reactivation in MDMs (*53*). Latently infected T-cells, monocytes, and macrophages were evaluated for levels of latency reactivation in the presence of a glutathione peroxidase (GPx) inhibitor (Tiopronin), a Trx inhibitor (PX12), and a TrxR inhibitor (Auranofin). A non-redox related LPA reported in T-cells (D106; NSC 155703) acted as a negative control (*18*). In the case of T-cells (JLats) and monocytes (TLats), the drugs were added in combination with TNF-*α* for 24 and latency reactivation was subsequently measured by flow cytometry. All four compounds suppressed HIV reactivation with varying degrees (Fig. 5A). In T-cells, PX12 and Auranofin were the strongest suppressors of HIV reactivation, and in monocytes, PX12 was the strongest suppressor of reactivation. For MLats, the compounds were added after PMA removal (day 0 post-differentiation) and kept in the media until analysis. Contrary to results in T-cells and monocytes, both PX12 and Auranofin enhanced HIV latency reactivation in MLats by ∼3 fold after 2 days (Fig. 5A). On the other hand, both D106 and Tiopronin did not affect reactivation of MLats. This indicates inhibition of the Trx/TrxR system can affect the regulation of HIV latency in a cell type-dependent fashion.

**Figure 5.**
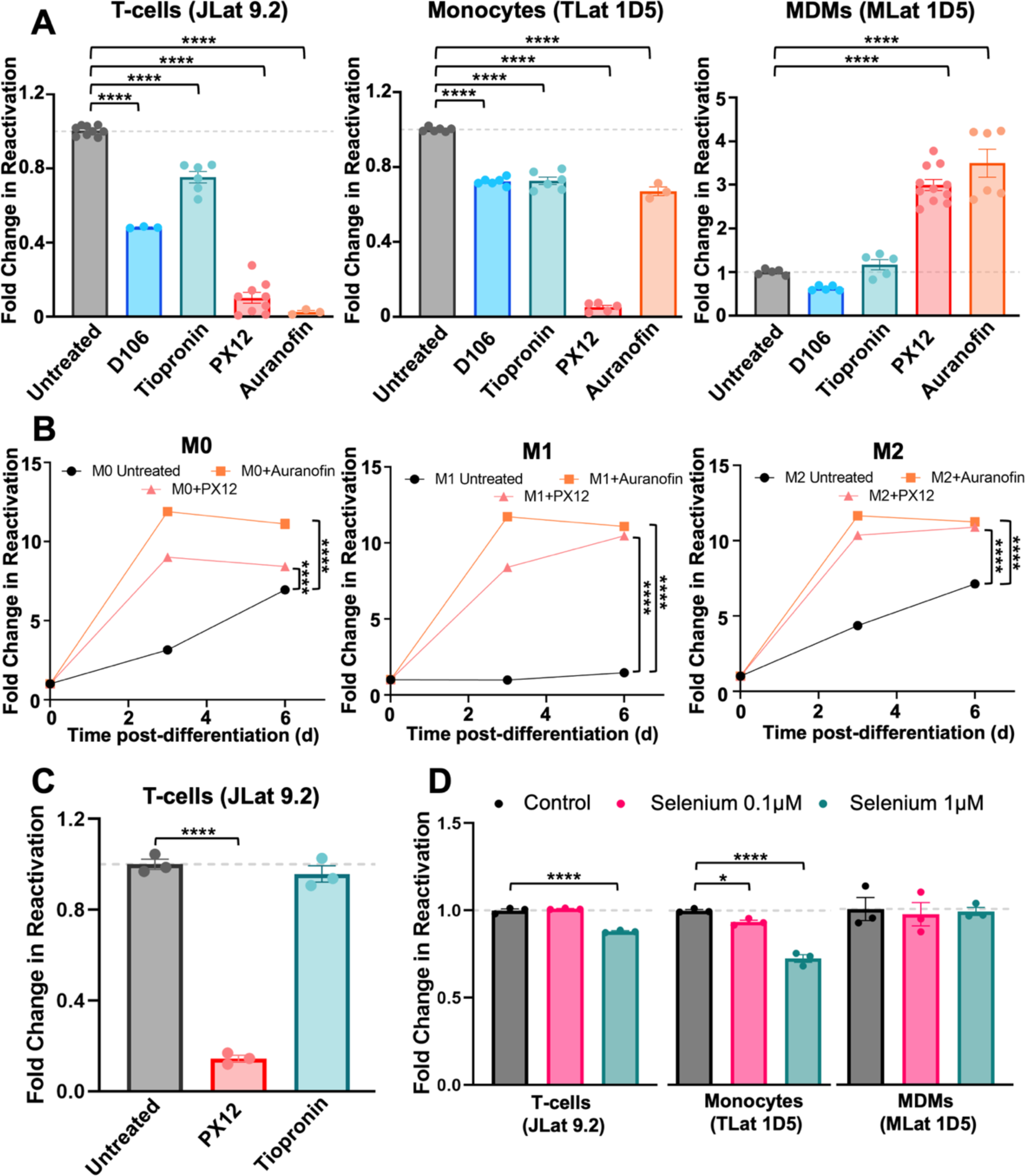
Macrophages differentially respond to modulation of redox homeostasis. (A) TNF alone or in combination with 10mM D106 (NSC 155703), 4mM Tiopronin, 60 µM PX12, or 4µM Auranofin were added to latently infected T-cells (Left) and monocytes (Middle) for 24h to challenge HIV reactivation. The same compounds were added to macrophages (Right) at day 0 post-differentiation for 48h to challenge HIV reactivation. Data points represent at least three replicates from 1-3 independent experiments ± SEM. (**B**) HIV reactivation was quantified for M0, M1, and M2 MLats in the presence of Auranofin and PX12 over time. Data points represent the mean of three independent replicates ± SEM. (**C**) PX12 and Tiopronin were added to latently infected T-cells in combination with PMA for 24h to assess whether there is an LRA-dependent response to redox protein inhibitors. Data points represent three independent replicates ± SEM. (**D**) Selenium was added to latently infected T-cells and monocytes for 72h, followed by TNF stimulation for 24h to achieve HIV reactivation. For MDMs, selenium was added at the time of differentiation and kept in the media for 72h. Data points represent three independent replicates ± SEM. Statistical significance was determined by performing a one-way ANOVA comparison with Dunnett correction (*: p<0.05, ****: p<0.0001).

To explore whether Trx/TrxR inhibition impacts HIV latency reactivation in the context of specialized macrophage function, we polarized MLats towards M1 and M2 phenotypes while treating with Auranofin and PX12. Addition of Auranofin and PX12 resulted in the enhancement of HIV reactivation across all macrophage subtypes (Fig. 5C), indicating similar mechanisms by which M0, M1, and M2 macrophages sense and respond to imbalances in cellular redox status. This suggests that regardless of the biological context and specialized function of latently infected MDMs, the modulation of ROS homeostasis by Trx/TrxR disruption promotes HIV transcription. We confirmed these observations to be independent of MLat clone (Fig. S5A), differentiation method (Fig. S5B), and LRA administered, as T-cells treated with PMA had a similar response to the compounds as T-cells treated with TNF-*α* (Fig. 5C). Further, we mimicked the T-cell and monocyte latency reversal assay for MDMs by adding TNF-*α* in combination with the compounds for 24h and obtained similar results (Fig. S5C). These results imply that there are cell type-specific differences in response to HIV latency modulation and that imbalances in cellular oxidation-reduction status can influence virus production.

Tiopronin and Auranofin target GPx and TrxR, respectively, selenium-containing enzymes that control cellular redox homeostasis. Previous reports have shown that selenium reduces HIV transcription by targeting the redox status of HIV Tat (*23*). Therefore, we wanted to explore whether selenium supplementation could affect HIV transcription in latently infected T-cells, monocytes, and MDMs. Selenium, in the form of sodium selenite, was added to T-cells and monocytes for 72h, followed by TNF-*α* stimulation for 24h to achieve latency reversal. In the case of MDMs, selenium was added with PMA during and after differentiation. Out of the two concentrations tested, selenium at 1μM suppressed HIV reactivation in both T-cells and monocytes (Fig. 5D). However, selenium supplementation did not influence HIV reactivation in MDMs, suggesting that additional biological mechanisms control HIV latency reactivation during differentiation. These findings further highlight the ability of cellular redox homeostasis to influence HIV transcription and suggest MDMs differentially regulate HIV latency compared to T-cells and monocytes.

### Inhibition of TrxR by Auranofin induces HIV expression by enhancing NF-kB and HIV promoter activity in MDMs

To investigate the mechanism of enhanced HIV latency reactivation in MDMs when cellular redox homeostasis is disrupted, we differentiated a clonal population of THP-1 monocytes integrated with the HIV LTR driving a destabilized d2GFP (LTR-d2GFP or LD2G) (*27*) into MDMs. We treated Ld2G monocytes and Ld2G MDMs with Auranofin and PX12 to track LTR promoter activity via flow cytometry. Compounds were added to monocytes in combination with TNF-*α* for 24h, and added to MDMs at day 0 post-differentiation, mimicking experimental conditions with TLats and MLats described above. Similar to results with JLats and TLats, Ld2G monocytes treated with Auranofin and PX12 showed decreased LTR activation, evidenced by decreased GFP expression (Fig. 6A). Ld2G MDMs treated with Auranofin showed increased LTR activation at both day 1 and day 2 post-differentiation, as observed with MLats (Fig. 6B). PX12, however, showed decreased LTR activity in Ld2G MDMs across both days, indicating its potent LRA-like effect in MLats is most likely driven through interactions with elements downstream of the promoter (e.g., viral proteins) absent in the Ld2G cell line.

**Figure 6.**
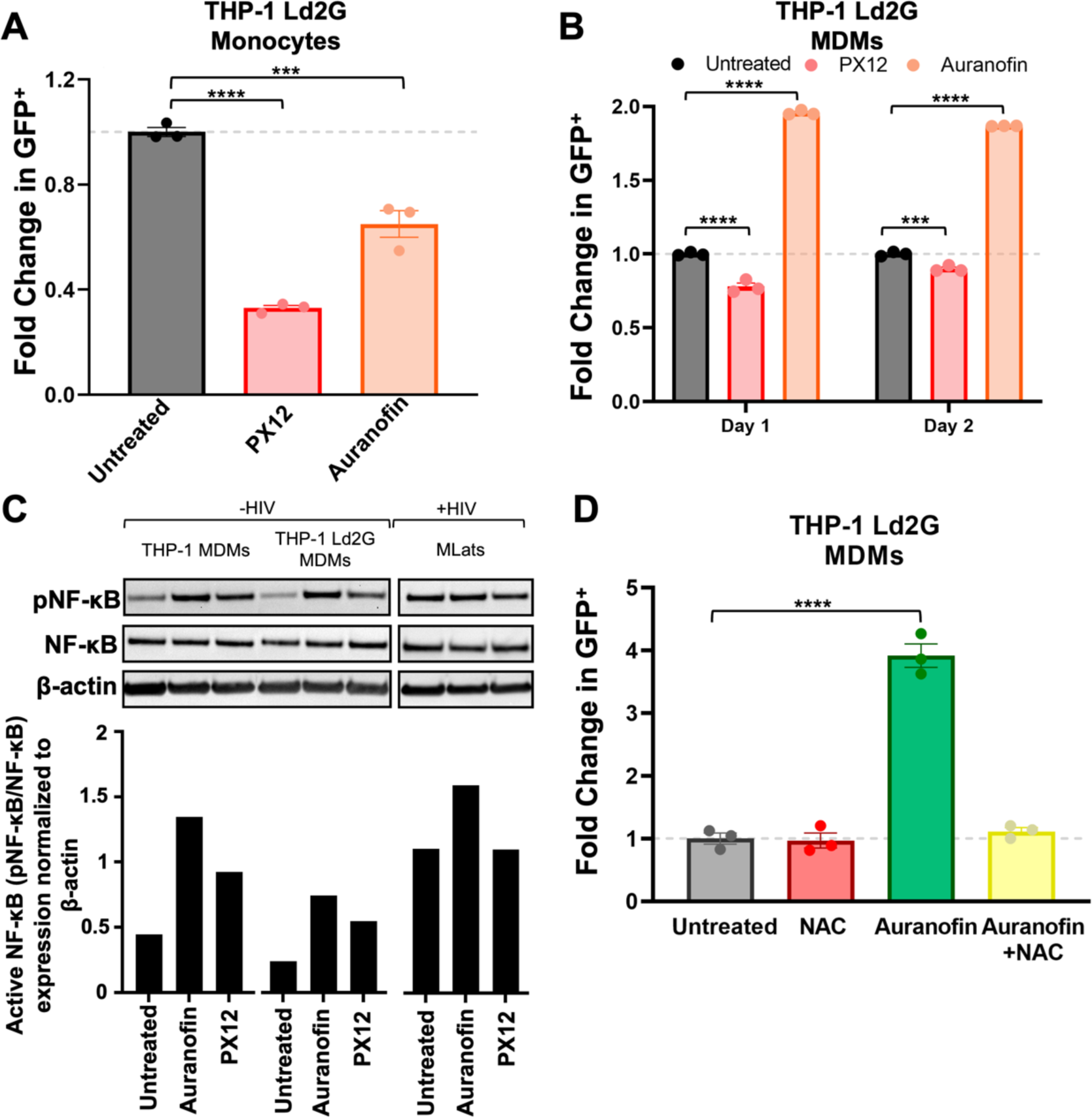
**Auranofin induces NF-κB activation of the HIV LTR promoter**. (A) TNF-a alone or in combination with 4µM Auranofin or 60µM PX12 was added to THP-1 Ld2G monocytes for 24h to evaluate LTR promoter activity. Statistical significance was determined by performing a one-way ANOVA comparison with Dunnett correction (***: p<0.001, ****: p<0.0001). (**B**) Auranofin or PX12 were added to THP-1 Ld2G MDMs at day 0 post-differentiation for 24h and 48h to evaluate LTR promoter activity. Statistical significance was determined by performing a two-way ANOVA comparison with Dunnett correction (***: p<0.001, ****: p<0.0001). (**C**) Western blot analysis of phosphorylated NF-κB (pNF-κB), total NF-κB, and B-actin (housekeeping) in THP-1 (uninfected) MDMs, THP-1 Ld2G (minimal HIV promoter circuit) MDMs, and MLats (latently infected) at day 1 post-differentiation (Top). Quantification of western blot results was performed by normalizing protein levels to B-actin. The ratio of pNF-κB to NF-κB was used to quantify activated levels of NF-κB, (Bottom). (**D**) N-acetyl-L-cysteine (NAC; 10µM) was added to THP-1 Ld2G MDMs at day 0 post-differentiation for 1.5h before the addition of Auranofin. LTR promoter activity was evaluated after 4h of treatment via flow cytometry. Statistical significance was determined by performing a one-way ANOVA comparison with Dunnett correction (****: p<0.0001). Data points represent three independent replicates ± SEM.

The enhancement of LTR activity and HIV reactivation by Auranofin in Ld2G MDMs and MLats, respectively, highlights a mechanism that might be largely dependent on the inducible DNA binding elements in the HIV LTR promoter. To assess specific promoter elements, we studied the key inducible transcription factor NF-κB (*2*) and evaluated active, phosphorylated pNF-κB (p65 subunit at Ser536), and total NF-κB levels in uninfected MDMs, Ld2G MDMs, and MLats treated with Auranofin and PX12 (Fig. 6C). Levels of pNF-κB in MLats were higher than in uninfected and Ld2G MDMs, which was expected as Tat feedback can stimulate NF-κB (*54*). Auranofin treatment induced pNF-κB to higher levels in uninfected MDMs, Ld2G MDMs, and MLats, being more pronounced in the former two (Fig. 6C). This indicates that Auranofin is capable of stimulating HIV transcription through induction of NF-κB activity. Interestingly, PX12 treatment also induced NF-κB activity in uninfected and Ld2G MDMs (to a lesser extent than Auranofin), but not in MLats, indicating alternative mechanisms by which it stimulates HIV transcription.

To dissect the specific mechanism by which Auranofin enhances NF-κB activity, we turned to the biological function of TrxR, the target of Auranofin. Inhibition of TrxR by Auranofin disrupts redox homeostasis and triggers oxidative stress, resulting in elevated levels of reactive oxygen species (ROS) (*20*). We hypothesized that elevated ROS levels in Auranofin-treated MDMs trigger the activation of NF- κB which subsequently causes HIV promoter activation. To test this, we treated Ld2G MDMs with N-acetyl-L-cysteine (NAC), a ROS scavenger and antioxidant, in combination with Auranofin, and evaluated LTR activity (Fig. 6D). NAC treatment suppressed the Auranofin-induced LTR activity back to untreated levels, highlighting ROS upregulation as the mechanism by which auranofin induces HIV expression in MDMs. Altogether, these results show that imbalances in cellular redox status from Trx/TrxR inhibition and elevated oxidative stress can trigger the expression of HIV in MDMs and potentially contribute to cell type-specific differences in HIV reactivation dynamics.

## Discussion

Here, we developed monocyte (TLat) and monocyte-to-macrophage differentiation (MLat) models of HIV latency to study the regulation of HIV latency within the myeloid reservoir. We purified single clones of latently infected monocytes to generate a library that exhibited broad levels of HIV reactivation capacity before and after differentiation (Fig. 1 and 2). These results recapitulate in myeloid cells the variability in HIV reactivation and responses to LRAs observed with several T-cell models of HIV latency (*55–57*). The differential regulation of HIV latency within these cellular reservoirs highlights the need for studying single cell populations of latently infected monocytes and MDMs when developing strategies to purge the latent HIV reservoir.

Differentiation of clonal populations of TLats into MLats revealed that monocyte-to-macrophage differentiation can trigger HIV reactivation, switching latently infected cells to virus producing cells (Fig. 2). We validated this finding in a physiologically relevant model using primary human CD14+ monocytes isolated from several donors, and showed that HIV reactivation longitudinally increased concomitant with macrophage differentiation, agreeing with a previous study reporting spontaneous and linear HIV reactivation over time in primary HIV-infected MDMs (*48*). Both our primary and cell line models showed heterogeneous levels of HIV reactivation, which could be driven by the proviral integration site and local chromatin environment (*29*). We showed two distinct biological mechanisms inherent to MDM differentiation that can trigger HIV reactivation from latency (Fig. 3). PKC signaling pathway activation induces functional changes that support biological function in MDMs (e.g. cell adhesion, cell cycle arrest, and cytoskeleton remodeling) (*58*, *59*). Activation of this pathway triggers HIV latency reversal through NF-κB signaling and HIV LTR activation (*60*). MDM differentiation also led to an increase in CycT1 protein levels which likely promote the escape from latency during differentiation through HIV Tat feedback (*10*, *11*, *61*, *62*). Given the transcriptome and kinome rewiring that occurs during differentiation, we propose that there are likely numerous pathways that support reactivation in latently infected MDMs. For instance, an RNA-seq study of uninfected MDMs revealed over 3,000 differentially expressed genes following differentiation, with PI3K/AKT, Notch, MAPK, and NF-κB representing significantly enriched pathways (*63*). These signaling pathways were previously utilized in “shock and kill” efforts for their ability to activate host transcription factors that induce HIV expression (*64–68*), indicating that inherent cellular monocyte-to-macrophage differentiation pathways are highly conducive to promoting HIV reactivation from latency.

Changes in macrophage polarization phenotype led to differences in the regulation of HIV latency (Fig. 4), with M1 polarization promoting latency and M2 polarization upregulating latency reactivation, agreeing with previous results in latently-infected primary MDMs (*48*). Contrastingly, HIV infection of M1 and M2 polarized macrophages restrict HIV replication, further highlighting differential regulation of HIV infection, replication, and transcription across MDM phenotypes and latency status (*45–48*). The suppressive effect exerted by M1 MDMs is thought to be partially mediated by the upregulation of HIV restriction factors, such as apolipoprotein B mRNA editing enzyme (APOBEC3A) impairing reverse transcription, and Tripartite Motif 22 (TRIM22) and Class II Trans-activator (CIITA) inhibiting proviral transcription. IL-4 stimulation of MDMs is thought to drive HIV reactivation in the M2 phenotype, although additional studies will help determine the specific factors contributing to the differential regulation of latency across macrophage phenotypes (*48*, *69*). Interestingly, MLats exhibited significant morphological differences from uninfected MDMs, acquiring more circular and compact shapes across all polarization phenotypes. Previous studies also showed that HIV induces morphological changes in macrophages to alter migration and promote virus spread (*70*, *71*). Future studies will be required to determine specific factors governing HIV latency reactivation and how morphological deregulation induced by HIV impacts cellular function across macrophage phenotypes (*48*, *69*).

We tested a panel of compounds recently reported to be LPAs in latently-infected T-cells and discovered latently-infected MDMs showed a differential response to treatment compared to T-cells and monocytes (*18*). Auranofin and PX12, inhibitors of the TrxR/Trx redox system, revealed a cell-type specific role in controlling HIV reactivation, acting as LPAs in T-cells and monocytes, and LRAs in MDMs (Fig. 5). This previously unobserved phenomenon demonstrates that the success of therapeutic candidates for “block and lock” or “shock and kill” could be hampered by the cellular diversity of the latent reservoir. Auranofin, an FDA-approved compound commonly used for the treatment of rheumatoid arthritis, was previously investigated in clinical trials for potential therapeutic applications in diseases such as cancer (*72–74*) and HIV (*25*, *75*). It was previously shown to decrease the cell-associated viral DNA reservoir in peripheral blood of macaques infected with SIVmac251 (HIV-1 homologue simian immunodeficiency virus) when used in combination with ART (*26*). However, most studies investigating the clinical potential of Auranofin for HIV focused on T-cells, and as shown here, this drug can exert differential effects based on cell type and concentration (Fig. S5D), which could lead to tissue specific differences in HIV latency reactivation. Additionally, it has been demonstrated that TrxR targets the oxidation status of Tat in macrophages, keeping Tat in a reduced state and preventing binding to the trans-activation response element (TAR) (*23*). Cumulatively, our results support a dual mechanism by which redox protein imbalances can trigger HIV transcription in latently-infected MDMs: Inhibition of TrxR/Trx by Auranofin (1) causes the accumulation of ROS, triggering NF-κB activation and thus transcriptional initiation (Fig. 6), and (2) prevents redox targeting of Tat disulfide bonds, therefore maintaining Tat in an oxidized state where it can bind to TAR and stimulate HIV transcription (Fig. 7). Future studies will be required to assess the influence of these pathways on HIV latency reactivation in diverse cell types and in response to latency modulating agents.

**Figure 7.**
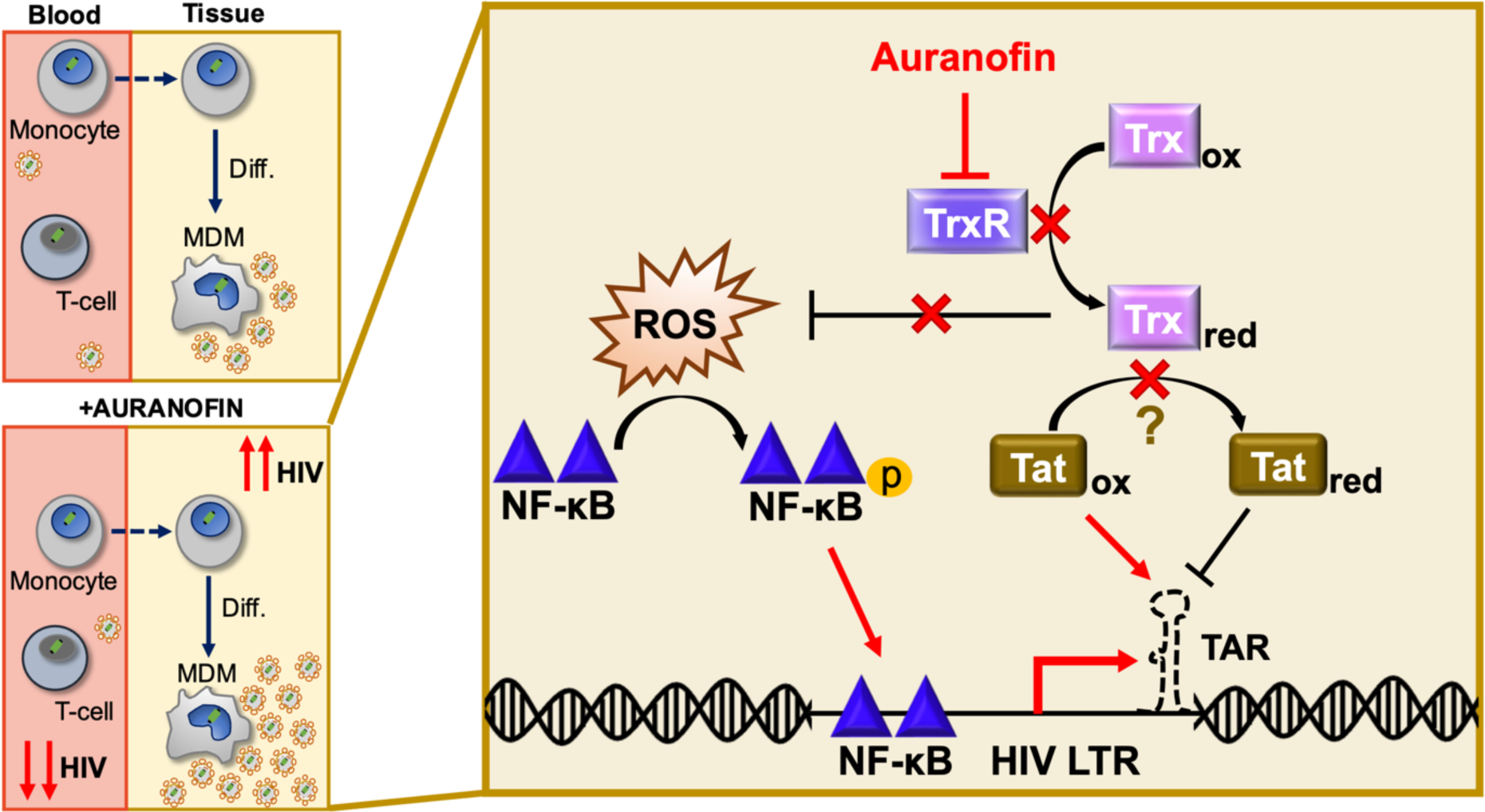
**Regulation of HIV latency in MDMs could present a risk to viral dissemination**. Monocyte-to-macrophage differentiation can induce HIV reactivation, potentially contributing to viral spread in tissues (Top left). The clinical candidate, Auranofin, reduces viral DNA in the blood (*25*, *26*) and promotes HIV latency in T-cells and monocytes, but induces HIV reactivation in MDMs (Bottom left). In MDMs, we propose TrxR inhibition by Auranofin leads to ROS accumulation, which induces NF-κB activity and activation of the HIV LTR promoter (Right). TrxR inhibition potentially diminishes substrate reduction, allowing Tat to remain predominantly oxidized, where it can bind to TAR and initiate HIV transcription (*23*).

Monocyte-to-macrophage differentiation and opposing cell type-specific responses to latency modulating agents present a new complication to HIV eradication strategies. These findings, together with recent evidence that MDMs harbor reactivatable latent reservoirs (*15*), highlight the potential threat that HIV infected monocytes and macrophages pose to viral dissemination and rebound during ART interruption in PLWH. Further, unintended consequences could arise from the administration of Auranofin in the clinic. Although able to reduce viral DNA in the blood, Auranofin could enhance latency reversal in tissue-resident macrophages, increase tissue viral burden and potentially lead to new infections in anatomical sanctuary sites with poor ART penetration. Thus, to achieve full control of the latent reservoir, latency clinical candidates will need to be thoroughly evaluated in all host cell types and biological contexts.

## Acknowledgments

We thank the staff of the Cytometry and Microscopy to Omics (CMtO) Facility at UIUC. JLat full length clones (9.2 and 15.4) were obtained through the NIH HIV Reagent Program from Dr. Eric Verdin. The NL4-3 ΔEnv EGFP vector was obtained through the NIH HIV Reagent Program from Dr. Robert Siliciano. The LTR-d2GFP vector was obtained from the Weinberger Laboratory at UCSF.

## Author contributions

Conceptualization: AB, RDD, CK

Methodology: AB, RAC, NA, KM

Investigation: AB, RAC, NA, KM

Visualization: AB

Supervision: RDD,CK

Writing—original draft: AB, CK

Writing—review & editing: AB, RDD, CK

## Data and materials availability

All data are available in the main text or the supplementary materials.

## Materials and Methods

### Cell culture

Jurkat T-cells (JLat 9.2) and THP-1 monocytes (ATCC) were cultured in RPMI 1640 media (ATCC) with L-glutamine and phenol red, supplemented with 10% fetal bovine serum (FBS; Gibco) and 1% penicillin/streptomycin (P/S; Gibco). Cells were maintained by diluting with fresh medium every 2-3 days. HEK293T cells were cultured in DMEM media (Corning) with 10% FBS and 1% P/S. THP-1 Ld2G cells were generated as previously described (*27*). Cells were incubated in 5% CO_2_ at 37 °C.

### Lentivirus production

The full-length HIV vectors NL4-3 ΔEnv EGFP (HIV Reagent Program) and HIV_GKO_ (Addgene plasmid #112234) were produced in HEK293T cells along with the VSV-G (Addgene plasmid # 8454) and ΔR8.2 (Addgene plasmid # 8455) plasmids for lentivirus generation. HEK293T cells (70% confluent) were transfected using FuGene 6 transfection reagent (Promega) according to the manufacturer’s instructions. Viral supernatant was collected after 24h and 48h, centrifuged at 500 x*g* for 5 min to remove remaining cells, and passed through a 0.45μm polyethersulfone membrane. Lentivirus was concentrated using the Lenti-X Concentrator (Takara Bio) according to the manufacturer’s protocol and titrated on HEK293T cells.

### Generation of a THP-1 monocyte HIV latency model (TLat)

A THP-1 monocyte HIV latency model was generated using the full-length vector NL4-3 ΔEnv EGFP as previously described (*27*). Lentivirus was generated by transfection of HEK293 cells as described above. Concentrated viral supernatant was used to infect THP-1 cells at a MOI < 1. Cells were allowed to grow and recover for ∼1 week following infection. GFP^-^ cells were sorted by FACS and cultured for an additional week. To reactivate latently infected cells, 10ng/mL TNF-*α* (R&D Systems) was added for 24 h, and GFP^+^ single-cells were sorted into 96-well plates (ThermoFisher Scientific). Cells were allowed to expand for over 3 weeks into clonal populations. To measure reactivation percentage of each clonal population, cells were stimulated with 10ng/mL TNF-*α* for 24 h. After TNF-*α* treatment, the number of GFP^+^ cells determined the reactivation percentage.

### Generation of a latent HIV-infected THP-1 monocyte-to-macrophage differentiation model (MLat)

Monocytes (naïve or latently infected) were differentiated to M0 macrophages using phorbol 12-myristate 13-acetate (PMA) (Cayman Chemicals). Cells at a density of 1x10^6^ cells/mL were incubated with 50ng/mL PMA for 24h. After 24h (day 0 post-differentiation), cells became adherent to tissue culture plastic and presented macrophage-like morphological characteristics. PMA was washed off and fresh PMA-free medium was added to the cells. At this point, MDMs can be kept in media for ∼1 week as M0 or polarized towards M1 or M2 phenotypes. Polarization was induced after a 24h rest period in PMA-free media (i.e., day 1 post-differentiation). MDMs were incubated with 20 ng/mL interferon gamma (IFN-y; Cayman Chemicals) and 250 ng/mL lipopolysaccharide (LPS; Millipore Sigma) (M1) or 30 ng/mL interleukin-4 (IL-4; Miltenyi Biotec) (M2) for 48h. Following the 48h incubation period (i.e., day 3 post-differentiation), polarization media was removed and fresh RPMI medium was added to the cells. MDMs (M0) were also generated by incubation with 100nM Vitamin D_3_ (VitD; Cayman Chemicals) or 1μM retinoic acid (RA; Cayman Chemicals) + 1μM VitD for 72h as previously described (*38*, *39*). For PKC activation studies, TLats were treated with the non-selective PKC inhibitor bisindolylmaleimide IX (BIM IX; Cayman Chemicals) for 30min prior to differentiation and kept in the media for the duration of differentiation. Cells were harvested at multiple time points for analysis (e.g., flow cytometry, western blot, qRT-PCR) by washing once with PBS (without Ca^2+^ and Mg^2+^), incubation with trypLE for 5 minutes to detach cells, and pelleting of cells at 300 x*g* for 5 min in a centrifuge.

### Generation of an HIV-infected primary monocyte-to-macrophage differentiation model

CD14+ monocytes were magnetically labeled with CD14 microbeads (Miltenyi Biotec) and separated from PBMCs (AllCells & STEMCELL Technologies) according to the manufacturer’s instructions. The resulting cell population was stained with CD3 and CD14 antibodies (ThermoFisher Scientific) to determine purity and evaluated via flow cytometry (Fig. S2A). CD14+ monocytes were incubated in RPMI media supplemented with 10% FBS, 1% P/S, and 50ng/ml M-CSF (R&D Systems) to induce macrophage differentiation. Media was changed every 2-3d. 7 days post-differentiation, cells were infected with HIV_GKO_ lentivirus at a MOI = 1 in the presence of 8μg/ml polybrene (Millipore Sigma) for 2h. Following infection, cells were rested in differentiation media and HIV infection was monitored by flow cytometry by evaluating GFP and mKO2 expression at 1, 2, and 3 days post-infection. Since EF1α was minimally expressed in infected primary cells compared to infected cell lines (Fig. S2B), we relied on GFP expression to monitor HIV infection and reactivation (*76*, *77*).

### HIV Latency Reversal Assays

JLat 9.2 and TLat 1D5 were diluted to 1x10^6^ cells/mL and treated with chemical modulators of latency. PX12, Tiopronin, and Auranofin (redox protein inhibitors) were acquired from Cayman Chemicals. D106 (NSC 155703) was acquired from the National Cancer Institute Developmental Therapeutics Program. PX12 (60μM), Tiopronin (4mM), Auranofin (250nM, 1μM, or 4μM) or D106 (10mM) were added to cells with and without TNF-*α* (10ng/mL) for 24h and analyzed via flow cytometry to determine HIV reactivation. In the case of JLat 9.2, PMA (200ng/ml) was also used as a latency reversal agent (LRA) to rule out an LRA-specific response to the inhibitors. In MDMs, cells were treated with the same redox protein inhibitors following differentiation and PMA removal from the media (day 0 post-differentiation). N-acetyl-L-cysteine (NAC; Cayman Chemicals) was added to MLats for 1.5h before the addition of Auranofin and kept in the media for the duration of treatment (3h). Cells were analyzed via flow cytometry at different time-points post-treatment.

### Protein detection by western blot

Cell pellets were washed in cold PBS and lysed with buffer containing 1% NP40 and protease inhibitor cocktail (Abcam). Whole cell lysates were obtained by centrifugation and removal of cell debris. Protein concentration of lysates was determined by Pierce BCA Assay (ThermoFisher Scientific). 15-20μg of total protein was loaded onto precast 4-12% iBlot2 Bis-Tris gels (ThermoFisher Scientific) and allowed to run for ∼90 minutes at 120V. Proteins were transferred onto a polyvinylidene difluoride (PDVF) membrane using the iBlot2 Dry Blotting System (ThermoFisher Scientific). The membrane was blocked in 5% non-fat milk or bovine serum albumin (BSA) for 1h and incubated overnight at 4°C with primary antibodies against Cyclin T1 (Cell Signaling Technologies), p24 (Abcam), phosphorylated NF-κB p65 (Ser536) (Cell Signaling Technologies), NF-κB (Cell Signaling Technologies), and B-actin (Cell Signaling Technologies) at dilutions recommended by the manufacturer. Proteins were detected by horseradish peroxidase-conjugated secondary antibody using SuperSignal West Pico PLUS chemiluminescent substrate (ThermoFisher Scientific) and imaged with an Invitrogen iBright FL1500 imaging system (ThermoFisher Scientific). Results were quantified by determining the area under peak for each band in ImageJ.

### Quantitative real-time PCR (qRT-PCR)

Total RNA was extracted from cells using the RNeasy mini kit (Qiagen) and concentrated with an RNA Clean & Concentrator kit (Zymo Research). First strand cDNA synthesis was performed using the Verso cDNA synthesis kit (ThermoFisher Scientific). The expression levels of Gag, EGFP, M1 & M2 markers (IL-6, TNF-α, IL-1β, TGM2, CCL22, and PPARγ) (*50*), GAPDH, and B-actin were detected by SYBR Green qRT-PCR (ThermoFisher Scientific) using an Applied Biosystems Real Time PCR system (ThermoFisher Scientific). The primers used were Gag forward 5’-CTGTCGACGCAGTCGGCTTGCT- 3’ & reverse 5’-GCTCTCGCACCCATCTCTCTCCTTCTAGCC-3’, EGFP forward 5’- CCCGACAACCACTACCTGAG-3’ & reverse 5’-GTCCATGCCGAGAGTGATCC-3’, IL-6 forward 5’-ACTCACCTCTTCAGAACGAATTG-3’ & reverse 5’-CCATCTTTGGAAGGTTCAGGTTG-3’, TNF-α forward 5’-CCTCTCTCTAATCAGCCCTCTG-3’ & reverse 5’- GAGGACCTGGGAGTAGATGAG-3’, IL-1β forward 5’-ATGATGGCTTATAGTGGCAA-3’ & reverse 5’-GTCGGAGATTCGTAGCTGGA-3’, TGM2 forward 5’-CGTGACCAACTACAACTCGG-3’ & reverse 5’-CATCCACGACTCCACCCAG-3’, CCL22 forward 5’-ATTACGTCCGTTACCGTCTG-3’ & reverse 5’-TAGGCTCTTCATTGGCTCAG-3’, PPARγ forward 5’- TACTGTCGGTTTCAGAAATGCC-3’ & reverse 5’-GTCAGCGGACTCTGGATTCAG-3’, GAPDH forward 5’-TGTACCACCAACTGCCTTAGC-3’ & reverse 5’-GGCATGGACTGTGGTCATGAG-3’, and B-actin forward 5’- GCGGGAAATCGTGCGTGACA-3’ & reverse 5’- AAGGAAGGCTGGAAGAGTGC-3’. Results were normalized using B-actin or GAPDH as a housekeeping gene and evaluated using the 2^-ΔΔCt^ method (*78*).

### Antibody staining and flow cytometry analysis

Flow Cytometry was performed using the LSR Fortessa with HTS or Symphony A1 instruments (BD Biosciences). Reactivation of latently infected cells was quantified by analyzing GFP fluorescence intensity and percentage of GFP^+^ cells. For phenotypic characterization of MDMs, cells were stained with an APC-conjugated CD11b antibody, a PE-conjugated CD80 antibody (M1), and a PE-Cy7-conjugated CD209 antibody (M2), a PE-conjugated HLA-ABC antibody (Class I MHC), an APC-conjugated HLA-DR antibody (Class II MHC), and an APC-conjugated CD4 antibody (ThermoFisher Scientific) according to the manufacturer’s instructions. Briefly, 5 µL of antibody was added to cell samples at a final volume of 100 µL. The staining was done on ice for 30 min in the dark. Cells were washed at least once in FACS staining buffer (ThermoFisher Scientific) to remove unbound antibody. Cell viability was analyzed by adding 1:1000 propidium iodide (PI) or 4′,6-diamidino-2-phenylindole (DAPI) prior to analysis. Apoptosis was quantified using an APC-conjugated Annexin V stain (BD Biosciences). Data was analyzed using FCS Express 6 software and gated for singlet live cells.

### Epifluorescence Microscopy and Analysis

M0, M1, and M2 THP-1 and TLat 1D5 MDMs were differentiated as described above and seeded directly in a 6-well glass bottom plate (ThermoFisher Scientific) at a density of 0.6x106 cells/mL. At day 0 (i.e., 24h post-differentiation), differentiation media was removed, wells were washed with warm PBS and fresh RPMI media was added. Live cell images were acquired in 6-well glass bottom plates on a Revolve R4 Epi-fluorescence Microscope (Discover Echo), equipped with a 5 MP CMOS Monochrome Camera (Fluorescence), using a 20x/0.25 NA Plan Achromat Air lens and 60x/1.25 NA Plan Achromat Oil lens (Olympus). GFP was detected with a mercury-free light-emitting diode (LED) light cube in the fluorescein isothiocyanate (FITC) channel using a 470/40 excitation filter, 525/50 emission filter, and 495 nm dichroic mirror. Cells were also imaged in the transmitted light channel. 2048 x 1536 images were acquired (pixel size: 3.45μm) at different time points (day 0, day 3, and day 6) with 40% laser power for 20x, 30% laser power for 60x, and an exposure time of 20-35 ms. Fiji/ImageJ was used to define regions of interest (ROIs) based on phase contrast outlines of cells for quantification of cell number, morphology, and GFP intensity.

### Statistical Analysis

All statistical analyses were performed using GraphPad Prism.

