## Supplemental Information for "Monocyte to macrophage differentiation and changes in cellular redox homeostasis promote cell type-specific HIV latency reactivation"

Blanco *et al.*

A

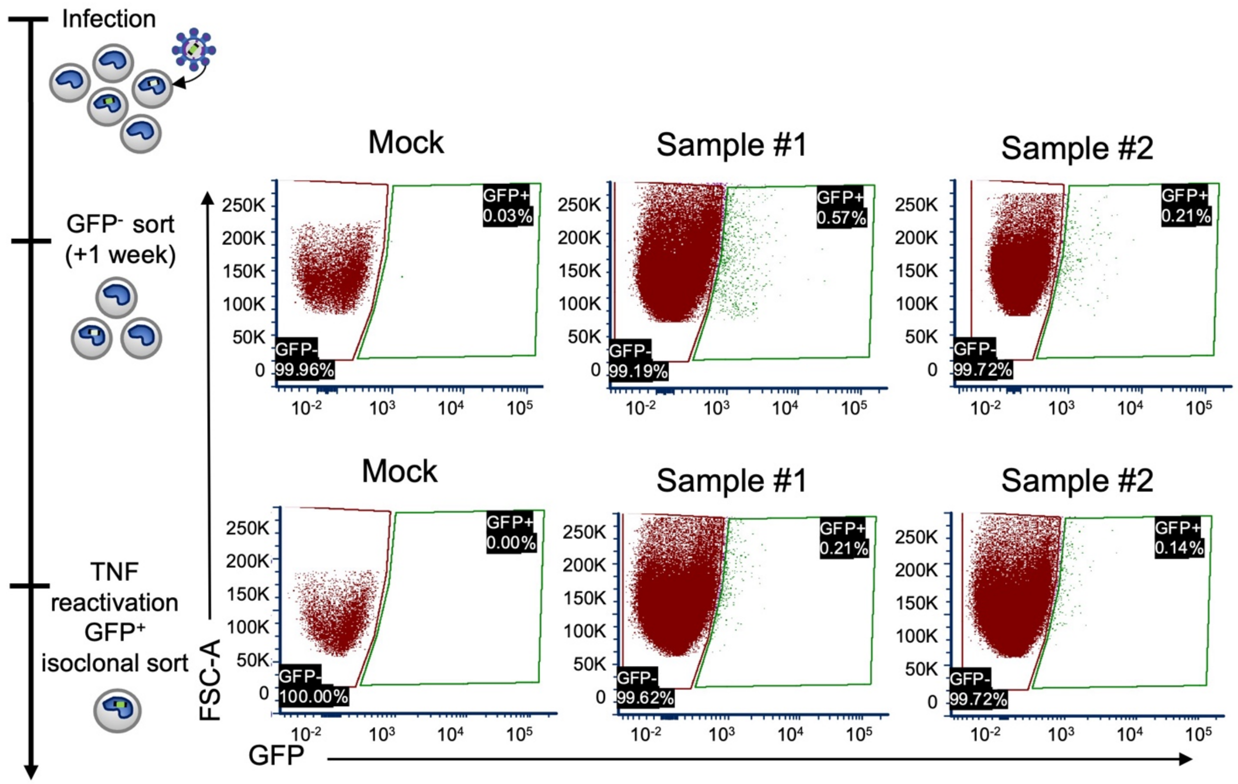


B

| Reactivation potential | # Clones | Mean unstimulated (basal) | Mean stimulated (+TNF-α) | Mean fold change in Reactivation |
|---|---|---|---|---|
| Low | 37 | 0.43 ± 0.36 | 0.84 ± 0.58 | 2.60 ± 2.29 |
| Intermediate | 25 | 1.01 ± 0.86 | 3.55 ± 1.99 | 4.67 ± 3.38 |
| High | 5 | 4.55 ± 5.28 | 57.95 ± 17.28 | 21.93 ± 12.54 |
| Low + Intermediate + High | 67 | 0.95 ± 1.78 | 6.11 ± 15.53 | 4.81 ± 6.44 |

**Figure S1.**

(**A**) Representative FACS plots showing the percentage of cells sorted at each stage of the TLat generation. Cells were infected at a MOI <1 to ensure a single integration per infected cell. Following infection, actively expressing NL4-3 ΔEnv EGFP cells comprised <1% of the entire population. TNF stimulation of GFP- cells resulted in <1% of HIV reactivation. GFP- and GFP+ gates were established based on mock-infected cells and applied to infected samples. (**B**) Quantification summary of the TLat library. Data represents mean ± SEM.

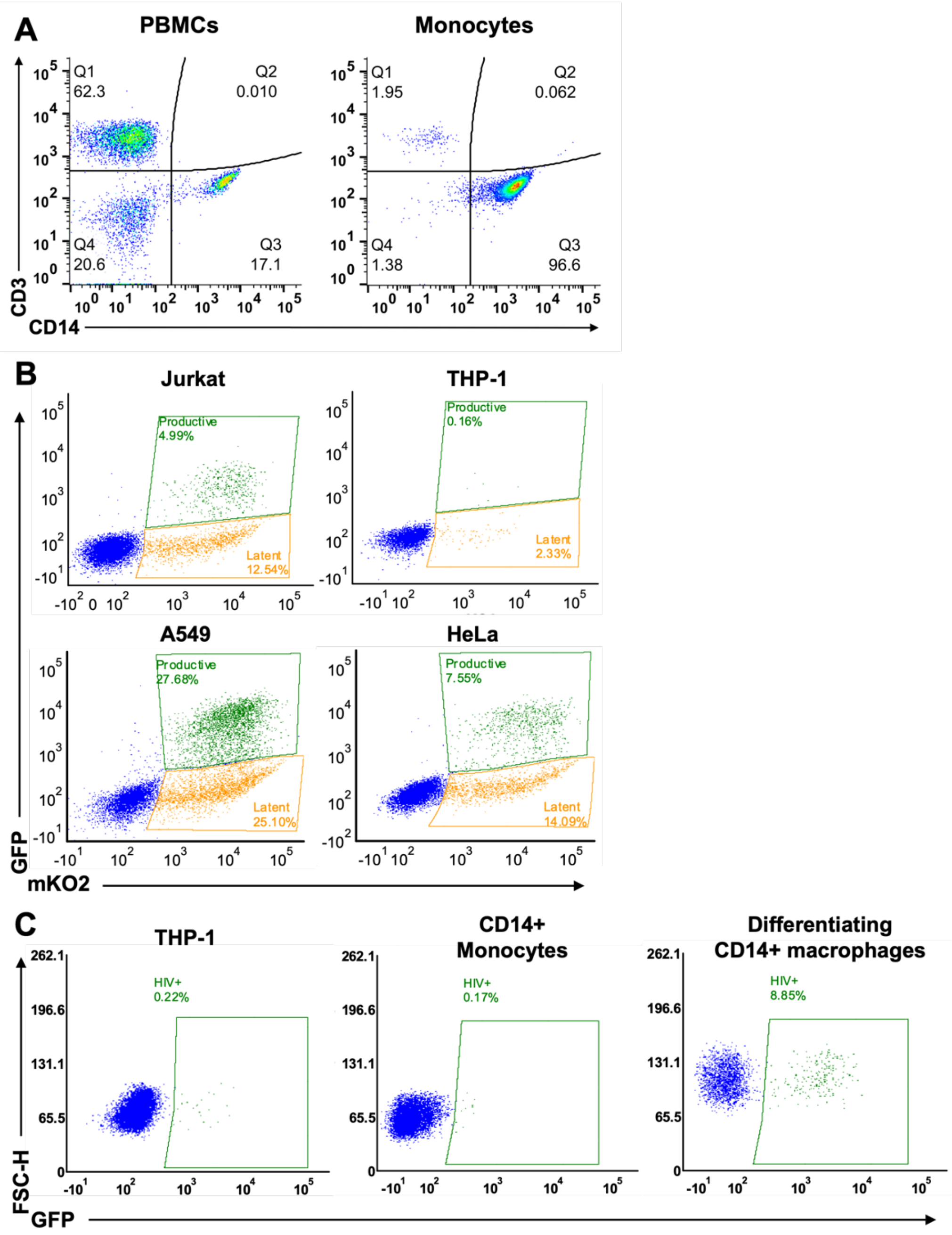


**Figure S2.**
(**A**) Representative flow cytometry plots showing the proportion of CD3+ and CD14+ cells in PBMCs before and after CD14+ monocyte separation. A minimal percentage of CD3+ cells is indicative of a pure CD14+ monocyte population. (**B**) Representative flow cytometry plots showing $HIV_{GKO}$ infection in multiple cell lines gives rise to two distinct populations: a mKO2+ GFP- population representing latent infections, and a mKO2+ GFP+ population represent productive infections. (**C**) Representative flow cytometry plots showing HIV production in THP-1 monocytes, CD14+ monocytes, and differentiating CD14+ macrophages 5 days post infection. Low numbers of HIV+ events in CD14+ monocytes limited confidence in interpreting the significance of infectivity changes in those conditions.

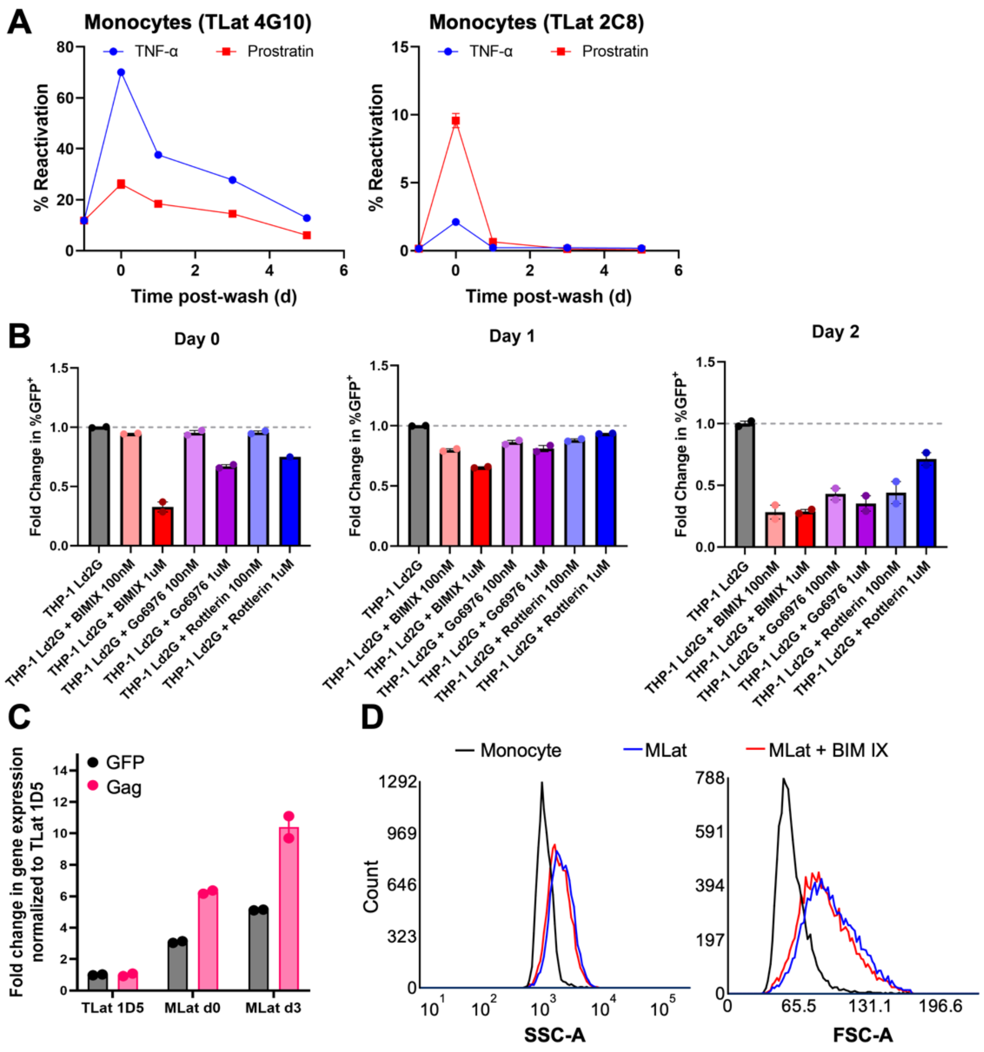


**Figure S3.**

(**A**) The PKC agonists (LRAs) TNF-$\alpha$, Prostratin, and PMA were added (at t=-1) for 24h to additional TLat clones (with intermediate and high reactivation profiles) to evaluate HIV reactivation following LRA removal (t=0). Data points represent at least two independent replicates ± SEM. (**B**) A clonal population of THP-1 monocytes integrated with the HIV LTR promoter driving a destabilized d2GFP reporter (LTR-d2GFP or LD2G) was treated with selective (Go6976 and Rottlerin) and non-selective (BIM IX) PKC inhibitors for 30 min prior to differentiation into macrophages and for the duration of the experiment. LTR transcription was quantified following differentiation by flow cytometry. Data points represent at least two independent replicates ± SEM. Dashed line indicates THP-1 Ld2G control. (**C**) mRNA levels of EGFP and Gag were analyzed by real-time quantitative PCR for uninfected THP-1 and HIV-infected (TLat 1D5) monocytes and MDMs. Results were normalized to uninfected THP-1s and B-actin mRNA levels. Data represents the mean of triplicates from two independent experiments ± SEM. (**D**) Representative flow cytometry side scatter (SSC-A) and forward scatter (FSC-A) histograms of monocytes, MLats, and MLats treated with BIM IX. Both side and forward scatter increased with differentiation and were not altered by BIM IX treatment.

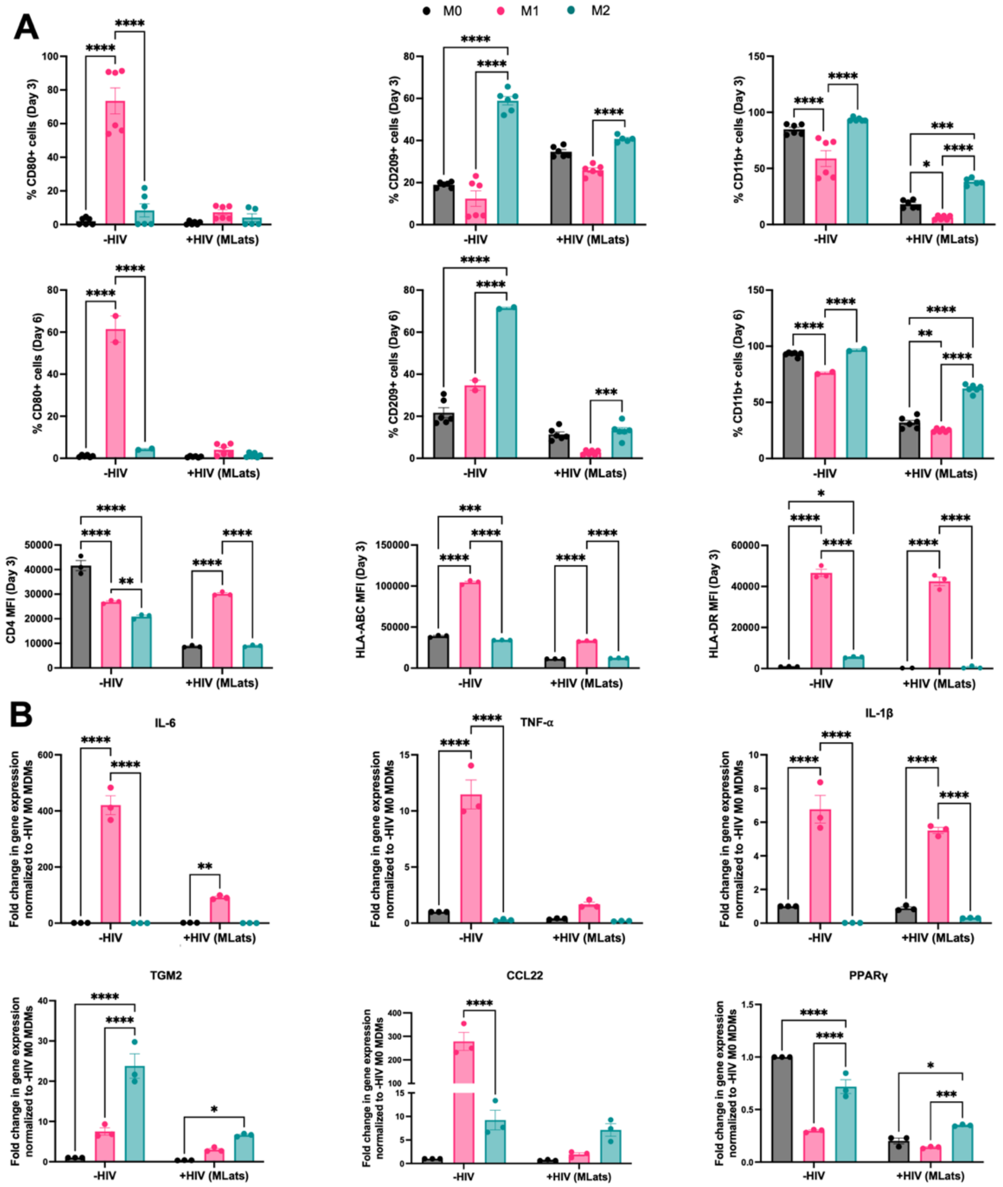


**Figure S4.**

(**A**) Flow cytometric analysis of uninfected or HIV-infected (MLat 1D5) MDMs staining positive for CD11B (general macrophage marker/M0), CD80 (M1), CD209 (M2), CD4, HLA-ABC (Class I MHC), and HLA-DR (Class II MHC) after differentiation. Latently infected MDMs showed decreased overall cell surface marker expression compared to THP-1 cells. Polarization of M0 MDMs to M1 and M2 phenotypes showed expected increases in CD80 and CD209 levels, respectively, which were less pronounced for infected cells.. (**B**) mRNA levels of M1 (IL-6, TNF-α, IL-1β) and M2 (TGM2, CCL22, PPARγ) cytokine expression in uninfected and HIV-infected (MLat 1D5) MDMs. Polarization of MDMs towards M1 and M2 phenotypes showed expected increases in M1-associated and M2-associated cytokine expression, respectively. Data points represent the mean of at least three independent replicates ± SEM. Statistical significance was determined by performing a two-way ANOVA comparison with Dunnett correction (*: $p<0.05$, **: $p<0.01$, ***: $p<0.001$, ****: $p<0.0001$).

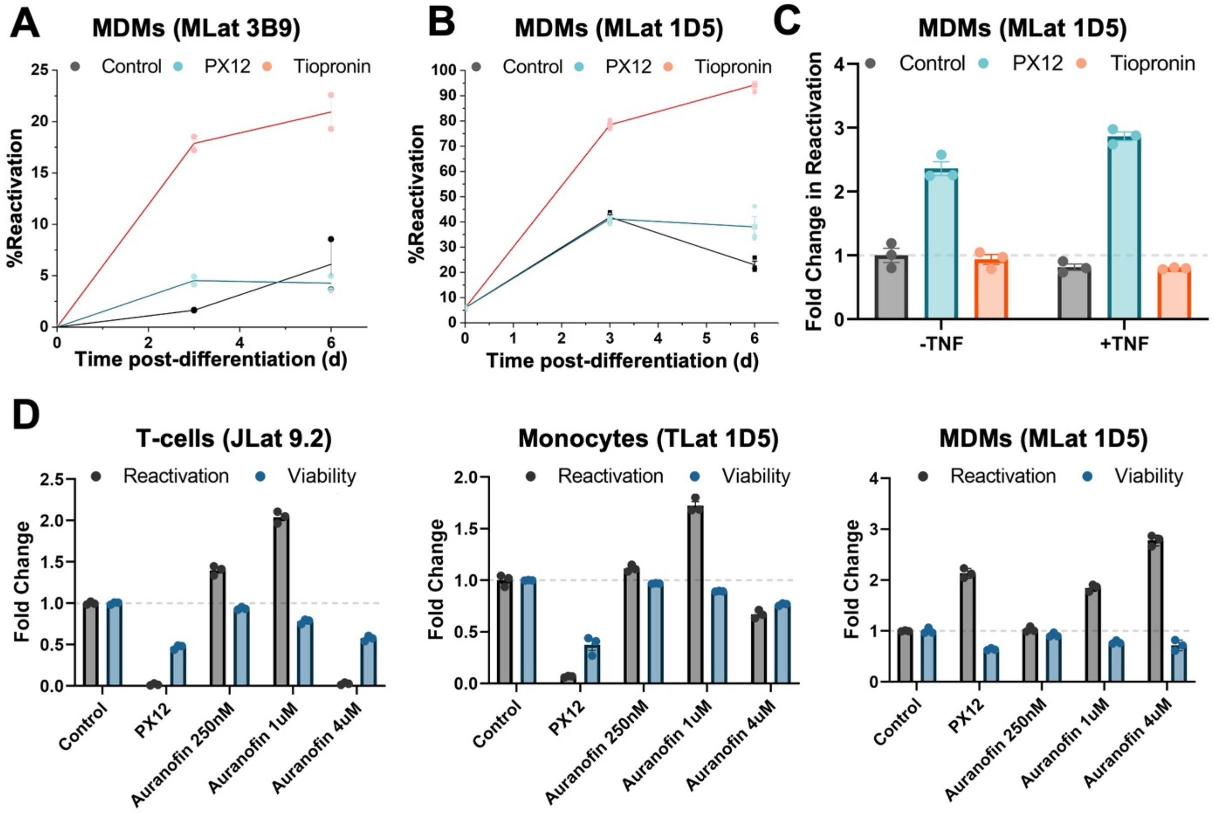


**Figure S5.**
(**A**) LPAs were added to an additional TLat clone with intermediate reactivation profile post-differentiation (MLat 3B9) to test reactivation levels. (**B**) Tiopronin and PX12 were added to MLat 1D5 generated by vitamin D3 to quantify HIV reactivation post-differentiation. (**C**) PX12 and Tiopronin were added to MLat 1D5 post-differentiation in combination with TNF for 24h to assess HIV reactivation. (**D**) Different concentrations of Auranofin were tested in T-cells, Monocytes, and MDMs. Reactivation and viability were quantified by flow cytometry. Data points represent at least two independent replicates ± SEM.

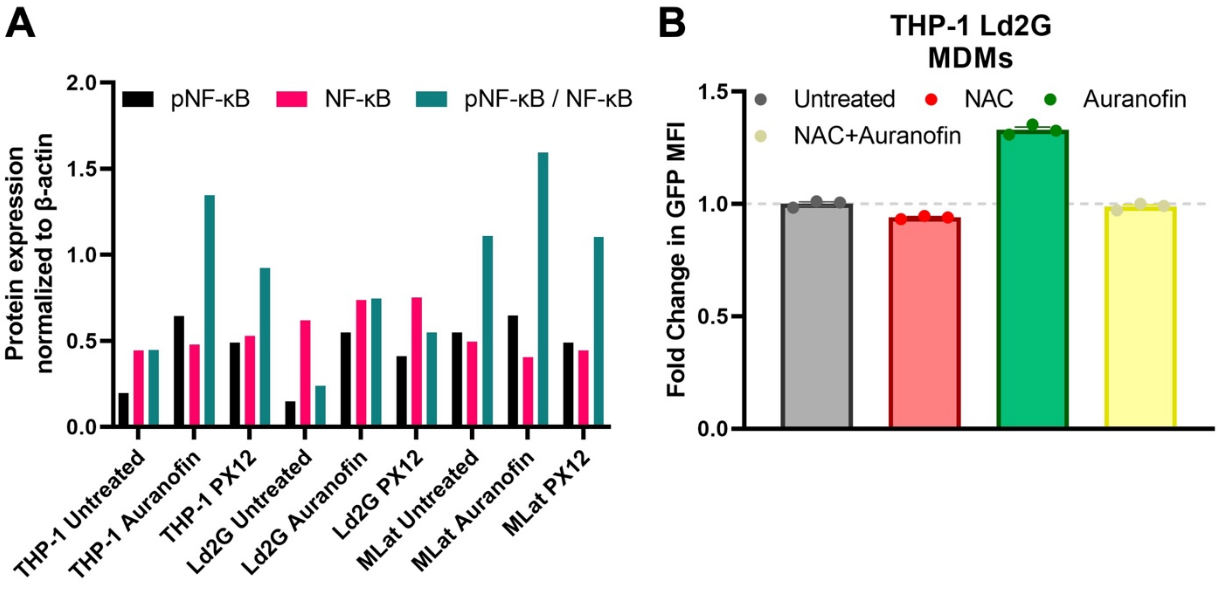


**Figure S6.**

(**A**) Phosphorylated NF-κB, total NF-κB, and active NF-κB (pNF-κB / NF-κB) protein levels normalized to B-actin. (**B**) Fold change in mean fluorescence intensity (MFI) of THP-1 Ld2G MDMs after 3h of treatment with Auranofin (4μM). N-acetyl-L-cysteine (NAC; 10μM) was added to THP-1 Ld2G MDMs at day 0 post-differentiation for 1.5h before the addition of Auranofin. Data points represent at least three independent replicates ± SEM.
